# Single cell RNA-seq identifies inflammation-induced loss of CFTR-expressing airway ionocytes in non-eosinophilic asthma

**DOI:** 10.1101/2022.04.26.489055

**Authors:** Ling Chen, Gabriela Araujo Hoefel, Prabuddha S. Pathinayake, Andrew Reid, Coady Kelly, Tan HuiYing, Richard Y Kim, Philip M Hansbro, Steven L Brody, Paul S Foster, Jay C Horvat, Carlos Riveros, Peter AB Wark, Gerard E Kaiko

## Abstract

Asthma is the most common chronic airways disease worldwide and the severe treatment resistant subtype of asthma is responsible for the majority of disease burden. Asthma is heterogeneous in nature and can be classified according to airway infiltrates as eosinophilic or non-eosinophilic (sometimes referred to as Type 2 low), which is further divided into paucigranulocytic (low levels of granulocytes), or neutrophilic asthma characterized by elevated neutrophils, and mixed Type 1 and Type 17 cytokines in airway tissue, sputum, and bronchoalveolar lavage. Severe non-eosinophilic asthma currently has fewer effective treatment options and many of these patients fail to qualify for newer biologic monoclonal therapies. The cystic fibrosis transmembrane conductance regulator (CFTR) is a key protein whose function is dysregulated in multiple respiratory diseases including cystic fibrosis and chronic obstructive pulmonary disease (COPD) and has proven a valuable therapeutic target. Using human bronchial epithelial cells (hBECs) isolated differentiated at air-liquid interface we demonstrated a reduced function of the CFTR in non-eosinophilic asthma. Characterization of the cell and molecular differences in airway epithelial cells between severe asthma subtypes using single cell RNA-sequencing (scRNAseq) revealed that airway epithelial cells from non-eosinophilic asthma, and in particular neutrophilic asthma patients, fail to differentiate into CFTR-expressing ionocytes compared with eosinophilic asthma or healthy donors. We identified a novel ionocyte transcriptional signature, which was present in both bronchial and tracheal airway epithelial samples indicating conserved anatomical gene regulation. Using protein markers and immunofluorescent quantification loss of ionocytes was confirmed in non-eosinophilic asthma hBECs. Similarly, ioncytes were also diminished in the airways of a murine model of neutrophilic-dominant but not eosinophilic allergen asthma models. Furthermore, treatment of hBECs from healthy donors with a neutrophilic asthma-like inflammatory cytokine mixture, but not IL-13, led to loss of ionocytes primarily due to IFN-γ. Inflammation-induced loss of CFTR-expressing ionocytes in airway cells from non-eosinophilic asthma may represent a key feature of disease pathogenesis and a novel drug target for this difficult-to-treat disease.

## Introduction

Severe asthma impacts about 1 in 10 people diagnosed with asthma, and is responsible for the majority of the healthcare burden both in terms of hospitalizations and costs of treatment (1). Severe asthma is heterogenous in nature consisting of patients with eosinophilic or Type 2 high asthma with classical eosinophilic airway inflammation (sputum or bronchoalveolar lavage (BAL)), with elevated Type 2 gene signatures (IL-4, -5, and -13), or patients with non-eosinophilic or Type 2 low airway inflammation (2). Non-eosinophilic asthma can be further sub-divided into neutrophilic asthma patients who demonstrate high airway neutrophilia and low eosinophils marked by elevated levels of Type 1 (IFN-γ, TNF-α) and Type 17 (IL-17A, IL-22) cytokines, or paucigranulocytic asthma patients with low levels of both eosinophils and neutrophils and a mixed inflammatory cytokine profile (3). Eosinophil levels are a key part of clinical decision-making and regulatory access to monoclonal antibodies targeting the Type 2 pathway. This has led to severe eosinophilic Type 2 high asthma now being successfully treated in the majority of patients, however non-eosinophilic asthma remains difficult-to-treat with few effective options (1). In particular, neutrophilic asthma has significant unmet need for new therapeutic options. Interestingly, this subtype of asthma bares many of the hallmark features of other chronic neutrophilic airway diseases such as chronic obstructive pulmonary disease (COPD), and cystic fibrosis (CF), including the airway neutrophilia, higher bacterial colonisation, mucus metaplasia and Type 1 and 17 dominated immune signatures (4–9). Given that COPD and CF have known defects in the CFTR (Cystic fibrosis transmembrane conductance regulator) chloride ion channel and that even mild defects in this pathway (in the case of adult-onset CF) are sufficient to induce the chronic airway disease, we hypothesized that there may be an induced dysfunction in this pathway in severe non-eosinophilic asthma. Furthermore, there is evidence that the risk of asthma is higher in non-CF individuals with CFTR gene variants (i.e. CFTR carrier heterozygosity) (10, 11). Additionally, a previous case series report also linked non-allergic asthma to abnormal nasal potential difference responses, a surrogate of CFTR function (12), suggesting that the disease state itself could induce changes in CFTR function.

In this study we demonstrate that airway epithelial cells cultured from non-eosinophilic asthma patients have reduced CFTR function and protein expression. To assess differences in subtypes of diseases, such as severe asthma, technologies with greater cellular resolution, such as single cell RNA-sequencing (scRNAseq), are required. Immune cell profiles in asthma have been studied using scRNAseq but the airway epithelial differences between asthma subtypes is yet to be investigated. Through scRNAseq we discovered that the diminished CFTR function is attributable to the loss of the ion channel-rich ionocyte in airway cells from non-eosinophilic asthma patients, in particular those with neutrophilic asthma. Ionocyte loss can be induced by inflammatory cytokines linked to neutrophilic airway disease including IFN-γ. Similarly these cells are lost *in vivo* in a murine model of neutrophilic, but not eosinophilic, asthma. Dysregulated ion channel transport through the CFTR and inflammation-induced loss of ionocytes may play a key role in the pathogenesis of non-eosinophilic asthma and represent a novel therapeutic target in this difficult-to-treat disease.

## Methods

### Participants

All subjects were 18 years or older, non-smokers or ex-smokers with less than 5 pack years smoked. Any subject demonstrating acute respiratory or any other illness within the previous 6 weeks was excluded. Spirometry was performed according to ATS/ERS guidelines (13). Exhaled nitric oxide was measured using a NIOX Vero (14). Healthy control subjects had no history of lung disease, were non-smokers with less than 5 packet year history of smoking and had normal spirometry. Asthma was defined by; a doctor diagnosis, together with a bronchodilator response of ≥12% and ≥200 mL of FEV1 or documented airway hyper responsiveness defined by a 15% decline in FEV1 during indirect bronchial provocation test respectively or evidence of demonstrable peak flow variability of more than 15% between the two highest and two lowest peak expiratory flow for 28 days. Asthma severity was defined by GINA guidelines, all participants required GINA step 4 treatment (15), with at least 500mcg of fluticasone propionate or equivalent daily along with a long-acting beta agonist. Approval of this study was obtained from Hunter New England Health human research ethics committee. Subjects and patients with severe asthma were recruited through the John Hunter Hospital clinics. Written informed consent was obtained from the study subjects.

### Human specimens

Bronchoalveolar lavage (BAL) fluid was processed, and differential cells counts were performed as previously described (16). Inflammatory asthma subtypes were defined based on BAL cell counts as previously reported (16); and defined as neutrophilic asthma when differential count was ≥70% neutrophils plus ≥0.99×10^6^/mL total cell count (TCC) and <3.5% eosinophils; eosinophilic asthma ≥3.5% eosinophils and <70% neutrophils, paucigranulocytic asthma when <3.5% eosinophils and <70% neutrophils. The clinical characteristics of the study subjects are shown in Table 1. Human bronchial epithelial cells (hBECs) were collected during clinical bronchoscopy by endobronchial brushings of the 3^rd^-4^th^ branch as previously described (17). Human tracheal epithelial cells (hTECs) were isolated from excess surgical tissue of lungs donated for transplant at Washington University in St Louis as previously described (18) and exempted from regulation as human subject research by the Washington University in St Louis Institutional Review Board.

### Murine asthma models induced by OVA, *Chlamydia muridarum*, and *Alternaria alternata*

Mouse studies were performed in accordance with protocols approved by the Animal Ethics Committee of The University of Newcastle. Wild-type BALB/c mice were sensitized to OVA (ovalbumin 50μg, intraperitoneal [i.p.] injection, Sigma-Aldrich) in Rehydragel^®^ (1mg (Reheis) in 200μl PBS under isoflurane anaesthesia. Mice were subsequently challenged intranasally (i.n.) with OVA (10μg/50μL sterile saline) on d12-13 to induce airways disease. Cmu/OVA mice on d14 were administered *Chlamydia muridarum* (ATCC VR-123 [Cmu], 100 inclusion-forming units, in SPG (sucrose-phosphate-glutamate buffer pH 7.5) i.n., and challenged with OVA again on d33-34 to exacerbate the airways disease. The OVA group of mice did not receive Cmu on d14 and instead received SPG vehicle control. Another set of wild-type BALB/c mice were challenged i.n. with *Alternaria alternata* (Greer) 5μg in 50μL PBS, three times per week, for five weeks. Control mice received either phosphate buffered saline or SPG buffered saline alone. Lungs were fixed in 4% paraformaldehyde overnight, washed in 70% ethanol, and then embedded in paraffin blocks for sectioning. BAL was collected and quantified by differential counts as previously described (19).

### Air-liquid interface (ALI) culture of primary airway epithelial cells

Healthy and asthma human bronchial epithelial cells (hBECs) were cultured in BEGM (Lonza) supplemented with BSA 8 µg/mL, 1% fungizone and 2% penicillin/streptomycin in a 6-well plate. Cells were trypsinized and seeded at 7 x 10^4^ cells per well in 6.5mm transwell with 0.4 μm pore polyester membrane inserts. Cells were submerged in BEGM/DMEM with supplements rhEGF (10ng/mL) and retinoic acid (30ng/mL). ALI was established by removal of apical medium, and cells were cultured with basal BEGM/DMEM with supplements rhEGF (0.5ng/mL) and retinoic acid (30ng/mL). Cultures were maintained at ALI and experiments conducted at 30 days. For cytokine experiments, cells received a mixture of Type 1/Type 17 cytokines containing IFNγ (10 ng/mL), IL-22 (10 ng/mL), IL-17A (10 ng/mL) and TNFα (1 ng/mL), or these individual cytokines once per week for the final 2 weeks on d21 and d28. d21 and d28, or IL-13 (20ng/ml) for the final 3 weeks.

### Ussing Chamber analysis of ion transport in airway epithelia

Ion transport properties in ALI cultured bronchial epithelial cells were studied by EasyMount Ussing Chamber Systems (VCC MC8 multichannel voltage/current clamp, Physiologic Instruments, Inc.). In brief, the apical and basolateral sides of the epithelium were perfused continuously with KRB buffer solution of the following composition: 115mM NaCl, 2.4mM K2HPO4, 0.4mM KH2PO4, 1.2mM CaCl2 dihydrate, 1.2mM MgCl2 hexahydrate, 25mM NaHCO3, 10mM HEPES and 10mM glucose, pH 7.4. The chambers were continuously gassed with 95% O2 – 5% CO2 and maintained at 37°C. The short-circuit current (Isc) and transepithelial resistance (Rt) were determined under voltage-clamp conditions. The Isc is a direct measure for the net movement of ions across the epithelium and recorded every 30 seconds. Data were acquired using Acquire and Analyze (version 2.3) software (Physiologic Instruments). The following specific compounds were added to the apical (A) and/or basolateral (B) bathing solutions in a standardized sequence. Cell membranes were equilibrated for 20 minutes in the presence of indomethacin (100µM, A+B) to inhibit endogenous chloride secretion caused by prostaglandin synthesis. Amiloride (100µM, A) was added to block amiloride-sensitive sodium ion absorption. Forskolin (10µM, A+B) and 3-isobutyl-1-methylxanthine (IBMX) (100µM, A+B) were added to activate cAMP-dependent CFTR chloride secretion. Carbachol (100µM, B) was added to stimulate cholinergic calcium ion transport. CFTRinh-172 (25µM, A+B) was added to inhibit CFTR-dependent chloride secretion. Isc responses to forskolin and IBMX (ΔIsc-Fsk+IBMX) was used as an indicator of maximum CFTR function.

### Single-cell RNA sequencing (scRNAseq) on airway epithelial cells

Airway epithelial cells were trypsanized from transwells monitored under microscopy and run through the 10XGenomics Chromium Next GEM v3.1 mRNA 3’ protocol as per manufacturer’s instructions. Briefly, single cells were partitioned into gel beads in emulsion with cell lysis and reverse transcription of RNA introduces cell barcoding. This is followed by PCR amplification, clean-up, and double-sided size selection with SPRIselect reagent (Beckman Coulter), and adaptor ligation and sample indexing by PCR. hTECs were assessed using the same protocol but with v3 chemistry. We aimed to recover a maximum of 10,000 cells per sample (∼16,000 cells loaded) for hBEC experiments and 3000 cells per sample for hTEC (∼4,800 cells loaded) experiments. hBEC samples were mixed at an equimolar ratio after indexing and sequenced with a Novaseq 6000 S1 flow cell followed by de-multiplexing, all hBEC samples were isolated and sequenced simultaneous avoiding need for batch correction. hTEC samples were sequenced separately.

### scRNAseq computational data analysis

Statistical hypothesis testing with the exception of the likelihood-ratio test (LRT), which is one-tailed, were two-tailed, and exact p-values are reported, except where below the threshold of numerical precision (2.22×10−16). Pre-processing of 3’ droplet-based scRNA-seq data demultiplexing, alignment to the hg38 transcriptome and UMI-collapsing were performed using the Cellranger toolkit (version 3.0.2, 10X Genomics). The CellRanger workflow performs the sample tag and UMI-demultiplexing, reference genome mapping, thresholding and low-quality filtering, clustering and preliminary differential expression analysis. Briefly, sequencer BCL files are demultiplexed to samples by bcl2fastq. Initial inspection and quality control was performed on the CellRanger output. Overall information on sequencing and mapping per sample is listed in Table 2. Subsequent data analysis was performed in R (v4.0.2, v4.1.2 (20) and Bioconductor Biobase v2.54 (21) using packages: Seurat v4.0.5 (22), sctransform v0.3.2 (23), uwot v0.1.11 (24, 25), Rtsne v0.15 (26, 27) fgsea (28), gage v2.44 (29).

The criteria used to detect a genes expressed in a cell is dynamically adjusted by CellRanger based on the overall profile of UMI counts per cell, the chemistry version used and the background read profile. For each cell, Cells with higher than 30% of total cell reads in mitochondrial genes or with less than 250 genes per cell detected were excluded from downstream analysis. In each sample, data normalisation was performed by a regularised negative binomial regression adjusted by the (residual) percentage of reads in mitochondrial genes, using the SCTransform function in Seurat (23). Then, PCA, UMAP and tSNE projections were performed and the 20 nearest neighbours on the 10 major principal components were computed. An iterative clustering was performed on the kNN data adjusting the resolution parameter to obtain a predetermined number of cell clusters on the dataset, using the simple modularity optimisation graph-based shared nearest neighbour method (30).

Differential gene expression analysis was performed on the regularised pseudo log count data using a Wilcoxon ranked sum analysis on each cell cluster versus all others, identifying genes overexpressed in the cluster of interest. Detected clusters were mapped to known cell types using known overexpressed gene markers (Table 3) using leading edge gene set enrichment analysis with an adjusted p value of 0.05 (28). The cluster cell type identification was then confirmed by verifying the assignment was the most significant enrichment assignment using GAGE (generally applicable gene set enrichment for pathway analysis) (29). During the preliminary data analysis due to the large proportion of basal and club cells, multiple clusters expressed high levels of markers for these two cell subtypes. This led to the iterative clustering procedure, reducing the resolution to obtain a more meaningful set of cell subtype clusters. Cell subtype signature genes used in the Venn diagram analysis to identify common signatures that were then used in pathway analysis were obtained by using a minimum FDR of 0.05 for adjusted p values. Genes were then ranked by these adjusted p values, and the top 100 ranked genes for each cluster/cell subtype was used in downstream analysis.

### Pathway analyses

Multiple pathway analyses were preformed using the genelists derived from scRNAseq analysis as described. This included Ingenuity Pathways Analysis (IPA), and use of ConsensusPathDB and ENRICHR (31, 32) for Gene Ontology (GO) for biological and molecular process, multi-pathway over-representation statistical analysis, and Human Phenotype Ontology. Library of Integrated Network-Based Cellular Signatures (LINCS) L1000 dataset was used to investigate ligands known to regulate gene signatures similar to the ionocyte gene signature.

### Immunofluorescence

Transwell membranes containing hBECs at the end of ALI cultures were fixed using 4% pPFA for 15 min at room temperature. For whole mount staining membranes or embedded in HistoGel (Thermo Scientific) as previously described (33). Sectioned membranes underwent heat-mediated antigen retrieval with sodium citrate buffer pH 6.0. Whole mount membranes or sections were blocked and permeabilized using PBS containing 3 % BSA, 5 % Donkey serum and 0.5 % Triton X-100 for 1 hour at room temperature. Additional blocking was applied using Background Sniper (Biocare Medical) for 10mins at room temperature. Cell membranes were incubated overnight at 4°C with the following primary antibodies anti-CFTR (1:100, 5µg/ml, R&D MAB23031), anti-FOXI1 (1:250, 2µg/ml, Abcam ab20454), and anti-ASCL3 (1:100, 1µg/ml, Sigma-Aldrich HPA027032), anti-Mucin 5AC (MUC5AC, 1:500, 2µg/ml, Abcam ab212636), anti-acetylated tubulin (Ac-tubulin, 1:1000, 1µg/ml, Sigma-Aldrich T7451), anti-p63 (1:50, 20µg/ml, Abcam ab735), and anti-uteroglobin (CCSP, 1:100, 10µg/ml, Thermo Fisher MA5-34625) at 4°C overnight. Alexa Fluor 555 or 488 conjugated secondary antibodies (1:500, 4µg/ml, ab150106, ab150129, ab150074) were applied in dark at room temperature for one hour, followed by mounting with Fluoromount-G with DAPI (Invitrogen). The number of ionocytes (ASCL3 and FOXI1 or CFTR and FOXI1 double positives) were quantified per high power field (HPF). The length of cilia (Tubulin) was quantified and presented as µm per mm basement membrane (BM). MUC5AC, p63, and CCSP positive cells were quantified and presented as number of cells per mm BM.

### Measurement and analysis of ciliary beat frequency (CBF)

Beating cilia of ALI cultured hBECs (frequency 3 to 18 Hz) were captured on a Nikon eclipse Ti2 microscope (Nikon, Japan) with Video Savant 4.0 software using a high-speed digital video recorder. High speed video was recorded at a rate of 300 frames per second and captured a minimum of 512 frames. Measurements of median cilia beat frequency were made using the ciliaFA plugin (34), which utilizes a specific fast Fourier transformation (FFT) algorithm to accurately measure ciliary activity across an entire field. CiliaFA was used together with free open-source ImageJ software (ImageJ, US) and Microsoft Excel to perform analyses.

### Statistics

Statistical analysis was performed by using GraphPad Prism software (Version 9.0). One-way ANOVA was used to identify differences between 2 or more experimental groups with correction for multiple comparisons. Student unpaired or paired *t* tests were used where appropriate for comparisons of 2 groups. All values are considered significant at a *P* value of less than 0.05.

## Results

### CFTR function is diminished in bronchial epithelial cells from severe non-eosinophilic asthma patients

Severe non-eosinophilic asthma patients have few treatment options. These patients are type 2 low and can also be subdivided into those that have high versus low airway neutrophils referred to as neutrophilic or paucigranulocytic asthma, respectively. The cellular and inflammatory features of this chronic airways disease and reports of abnormal nasal potential differences (12), bare similarities to other chronic respiratory diseases such as CF and COPD that have known genetic or acquired deficiencies in the CFTR. Given these cell and molecular similarities, we assessed whether CFTR function was altered in human bronchial epithelial cells (hBECs) from severe non-eosinophilic asthma patients. Patient cells were differentiated under standard conditions at air-liquid interface (ALI) and assessed by standard Ussing chamber ion transport measurements. Results revealed a significant reduction in CFTR function and chloride ion transport in non-eosinophilic asthma hBECs (Fig 1A). This was confirmed by inhibition of the CFTR channel with the small molecule CFTR-72 (Fig 1B). Amiloride stimulation, which inhibits the ENaC epithelial sodium ion channel showed a smaller reduction in ion movement (ENaC activity) in non-Type 2 asthma compared to healthy controls although this did not reach statistical significance (Fig 1C-D). To validate this data we quantified the number of CFTR protein expressing cells in the hBECs and confirmed that non-eosinophilic asthma has a significantly reduced number of CFTR+ cells versus healthy controls (Fig 1E). These data reveal a dysregulation of CFTR ion channel transport in differentiated airway epithelial cells from patients with non-eosinophilic asthma. Given the importance of CFTR to chloride ion movement, airway mucus dehydration and hypersecretion, and homeostatic lung defences through mucociliary clearance this finding may have implications for the pathogenesis of severe non-eosinophilic asthma.

**Figure 1.**
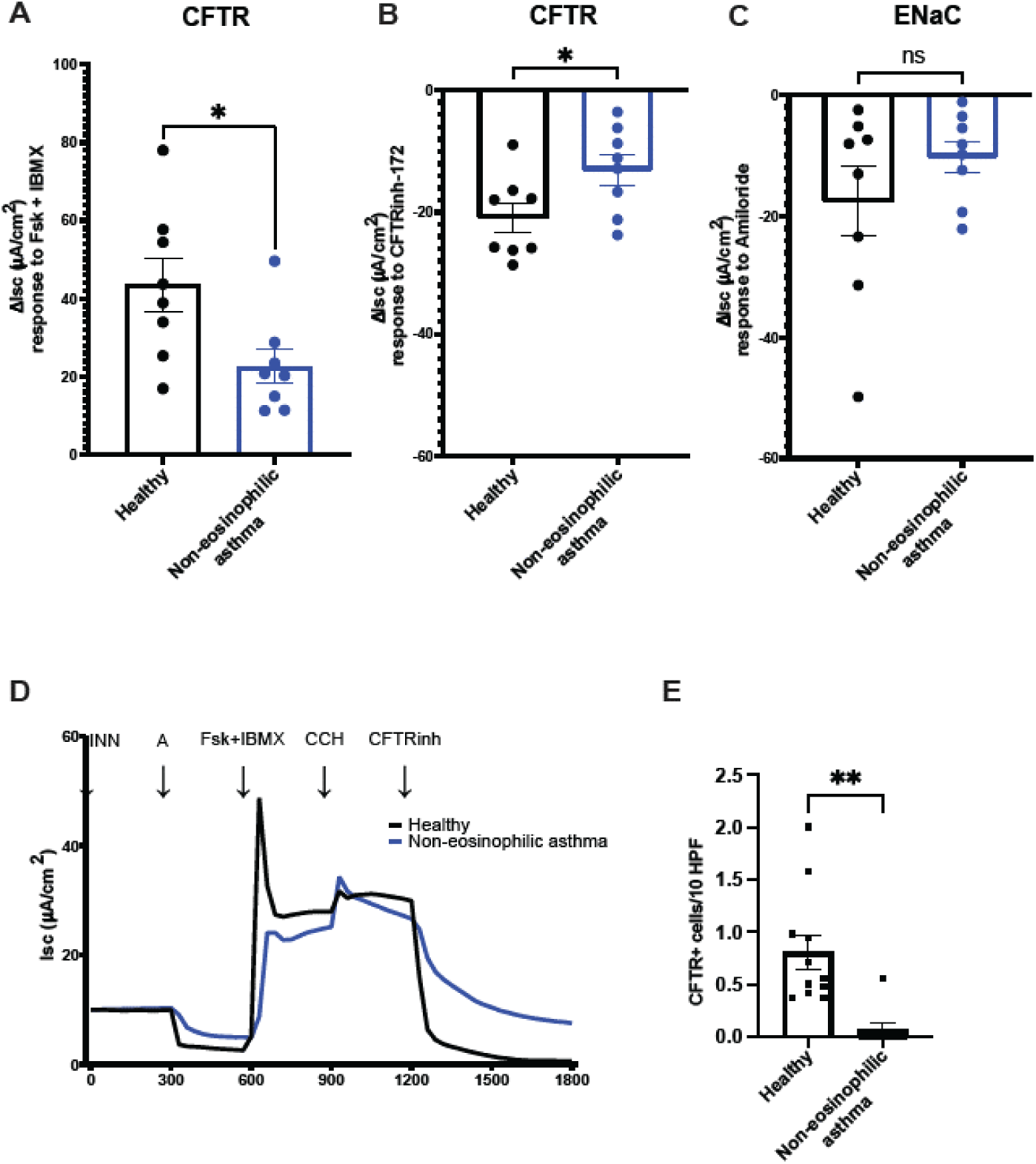
Ion transport measurements in healthy and non-eosinophilic asthma human bronchial epithelial cells (hBECs) at Air-Liquid Interface (ALI). The delta values of short circuit currents (ΔIsc) for (A) CFTR-Forskolin + 3-isobutyl-1-methylxanthine (IBMX) stimulated, (B) CFTRinh-172 inhibitor specifically blocking CFTR-dependent activity, and (C) Amiloride-sensitive ENaC currents for healthy (n=8) and non-T2 asthma (n=8) hBECs grown at ALI for 28 days. (D) Representative Isc responses of hBECs (circle line represents a healthy subject, and square line represents a non-eosinophilic asthma subject) after sequentially stimulated with amiloride (100 µM), forskolin (10 µM) + IBMX (100 µM), carbachol (100 µM), and CFTRinh-172 (25 µM). INN, indomethacin; A, amiloride; Fsk, forskolin; IBMX, 3-isobutyl-1-methylxanthine; CCH, carbachol; CFTRinh, CFTRinh-172 inhibitor. (E) CFTR protein expression was quantified by immunofluorescent staining in hBECs from healthy or non-eosinophilic asthma to determine the number of CFTR+ cells per 10 high powered fields (HPF). Each circle or square represents one individual subject. Error bars represent standard error of the mean (Mean ± SEM). Unpaired *t* test was used to determine statistical significance, * p<0.05, **p<0.01.

### Single cell RNA-seq reveals loss of CFTR-expressing ionocytes in bronchial epithelial cells from severe non-eosinophilic but not eosinophilic asthma

To assess airway epithelial cell subtype and transcriptional differences between healthy and severe asthma subtypes we conducted scRNA-seq analysis on patient-derived ALI differentiated hBECs from health donors, severe eosinophilic asthma, and severe non-eosinophilic asthma, including neutrophilic and paucigranulocytic patients (characterised by bronchoalveolar lavage (BAL) cell counts, n=4 patients each) (Supp Fig 1A). Cell subtypes were clustered using Seurat and known subtypes classified using classical airway epithelial markers with leadingEdge analysis (Table 3). Some clusters shared features of multiple subtypes and were therefore likely in a progenitor transitory stage and were thus labelled by predominant expression as *SERPINB+* progenitors, *LAMB3+* progenitors, or secretory progenitors. We observed several minor abundance differences in subsets between healthy donors and severe asthma subtypes, however, there were two very clear differences revealed by the scRNAseq analysis. Bronchial epithelial cells from neutrophilic asthma patients showed a marked increased abundance of mature goblet cells compared to more immature secretory progenitors present in the other patient groupings (Fig 2A-B and Supp Fig 1B-C). Furthermore, the most striking difference between patient samples was the complete loss of airway epithelial CFTR-expressing ionocytes in neutrophilic asthma patients (no ionocytes detected among 4 patients hBECs) (Fig 2C). Similarly, there was also a reduction of ionocyte abundance in paucigranulocytic samples compared to the higher levels detected in hBECs (∼2% of cells) from healthy donors and eosinophilic asthma. These differences were confirmed by unsupervised hierarchical clustering of ionocyte signature genes demonstrating the absence of these cells and their transcriptional marker genes in neutrophilic asthma (Fig 2D and Supp Fig 2A). These novel data reveal a reduction of ionocyte differentiation in non-eosinophilic asthma and provide a mechanism to explain the reduced CFTR and dysregulated ion channel function in differentiated hBECs from non-eosinophilic asthma (Fig 1). Given that defects in the CFTR protein alone are sufficient to induce chronic airway obstruction and neutrophilia in CF, this observation likely carries important clinical significance in severe asthma.

**Figure 2.**
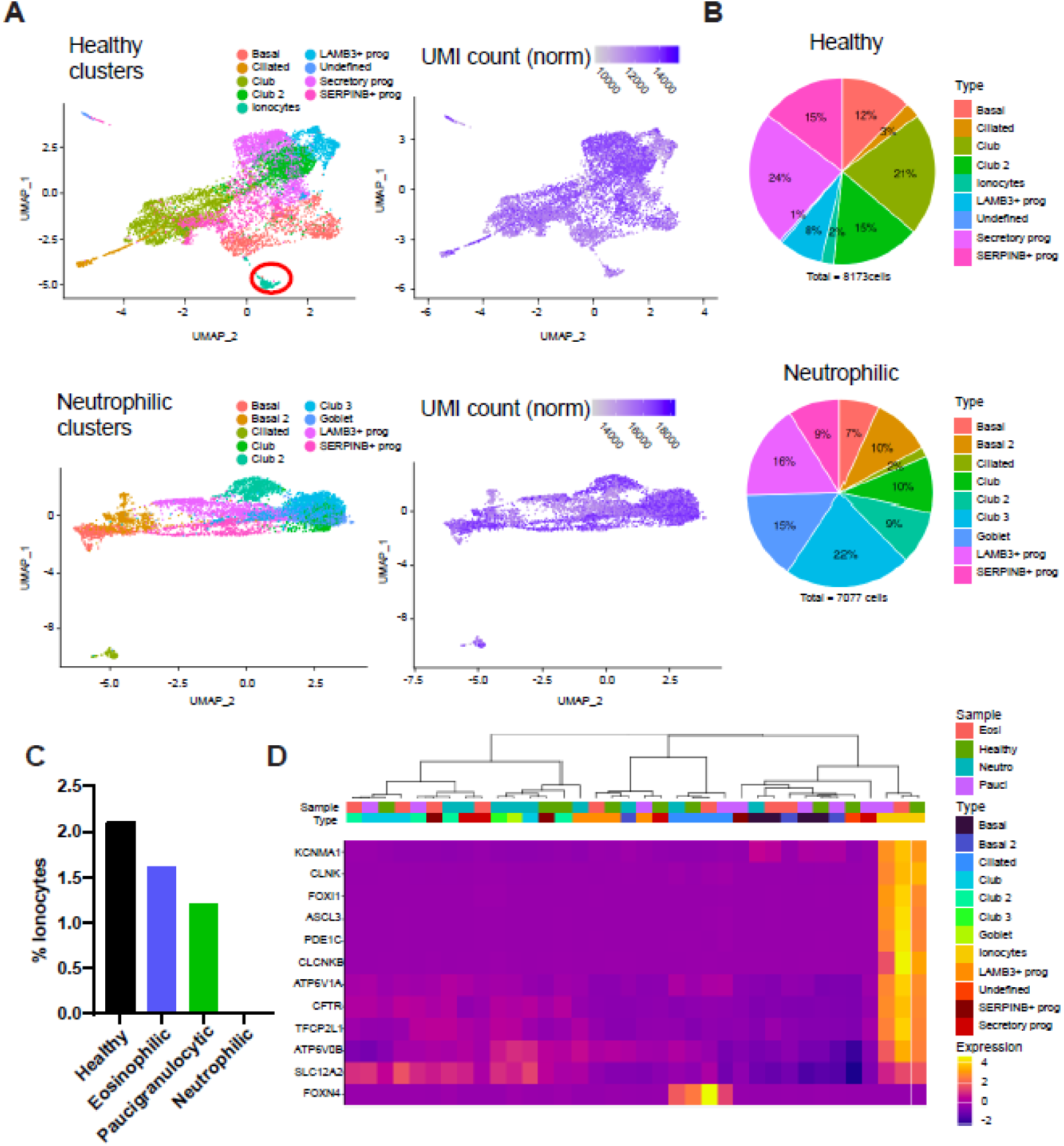
Single-cell RNA-seq analysis of hBECs. Single-cell RNA-seq was performed on bronchial epithelial cells at ALI generated from healthy subjects and neutrophilic asthma patients (n=4 each). (A) Cells were clustered by using a graph-based shared nearest neighbour method and plotted by UMAP together with heatmaps of gene UMI counts. 9 clusters of cells were identified in all samples and then known cell types were classified using classical gene markers enriched in the clusters through leadingEdge. Transitory progenitors or non-classified cell types were identified by predominant transcriptional signatures SERPINB+ prog, LAMB3+ prog, or Undefined. (B) The proportion of each cell subtype in healthy (8173 cells) and neutrophilic asthma (7077 cells) subjects are expressed (ionocytes in healthy = 2%, ionocyte absent in neutrophilic asthma). (C) Percentages of ionocytes in the healthy donors and patients with different types of severe asthma. (D) Unsupervised hierarchical clustering of all patient samples and cell subtypes (columns) by known ionocyte marker genes (rows), visualised using a heatmap plot.

### Ionocytes from trachea or bronchi possess a conserved gene signature with transcriptional machinery for chloride, iron, potassium, and pH regulation

Transcript expression of known ionocyte markers *FOXI1, ASCL3, PDE1C,* and *CFTR* in single cells was lost in hBECs from neutrophilic asthma (and reduced in paucigranulocytic asthma) compared to healthy donors or eosinophilic asthma (Fig 3A and Supp Fig 3). This analysis also identified the top 10 most differentially expressed genes in ionocytes from hBECs (*TMPRSS11E, ASCL3, CLNK, HEPACAM2, FOXI1, LINC01187, DGKI, STAP1, CLCNKB, PDE1C*) compared to all other epithelial cell subset clusters through pairwise comparison with every other cell cluster (Fig 3B and Supp Fig 4A-C). Similarly, this analysis identified key transcriptional markers of all other cell subsets. Through pairwise comparison in each sample for each cell cluster the top 100 marker genes characteristic of ionocytes in both healthy donors and severe eosinophilic asthma was identified and these 100 genes had 73% overlap (Fig 4A). These genes were enriched in pathways for ion channel transport including chloride, sodium, iron, and potassium ions, as well as pH regulation (Fig 4B and Supp Fig 5A-C). Human phenotype ontology analysis linked ionocytes and their gene expression to physiologic salt imbalances and metabolic pH dysregulation (Supp Fig 5C). To determine whether the ionocyte gene signature was conserved between different anatomical sites in the airways we conducted scRNAseq on human tracheal epithelial cells (hTECs) from healthy tissue similarly differentiated at ALI. Similar to bronchial cells we detected ionocytes in ∼1% of hTECs (Fig 4C-D and Supp Fig 5D). To compare the similarity between the top gene signatures in samples with clear ionocyte populations we examined the common marker genes in each of the healthy hTEC and hBEC samples as well as the hBECs from eosinophilic asthma. We identified a highly conserved ionocyte signature of 30 marker genes among these samples derived from different anatomical sites and 12 different patients (30% common between the 3 sets of patient samples) (Fig 4E). This signature included the characteristic marker genes *FOXI1, ASCL3,* and *CFTR* as well as novel marker linked to neuronal signalling. Pathway analysis showed strong enrichment for genes involved in ion channel transport (Fig 4E and Supp Fig 5E).

**Figure 3.**
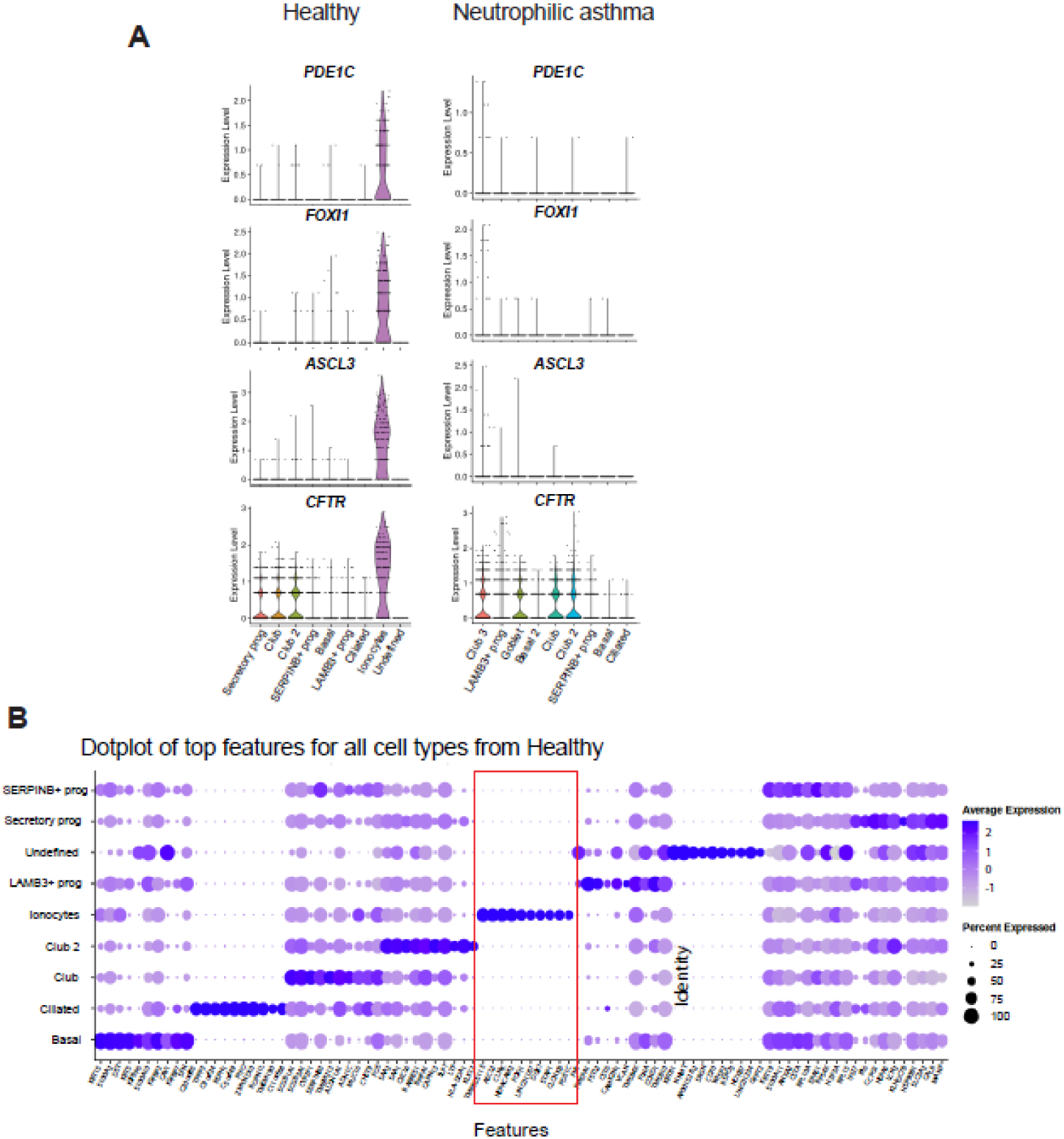
Identification of top ionocyte expressed genes in scRNAseq. (A) Violin plots of expression of ionocyte gene markers (*PDE1C, FOXI1, ASLC3* and *CFTR*) in each of the 9 cell subtype clusters from healthy subjects versus neutrophilic asthma patients. (B) Weighted dot plot showing top 10 features of each cluster from the scRNAseq data derived from cells of healthy subjects. Each dot is sized to represent the percent of cells in each cluster expressing the corresponding top 10 gene, and colors represent the average expression of each maker gene across within that cluster.

**Figure 4.**
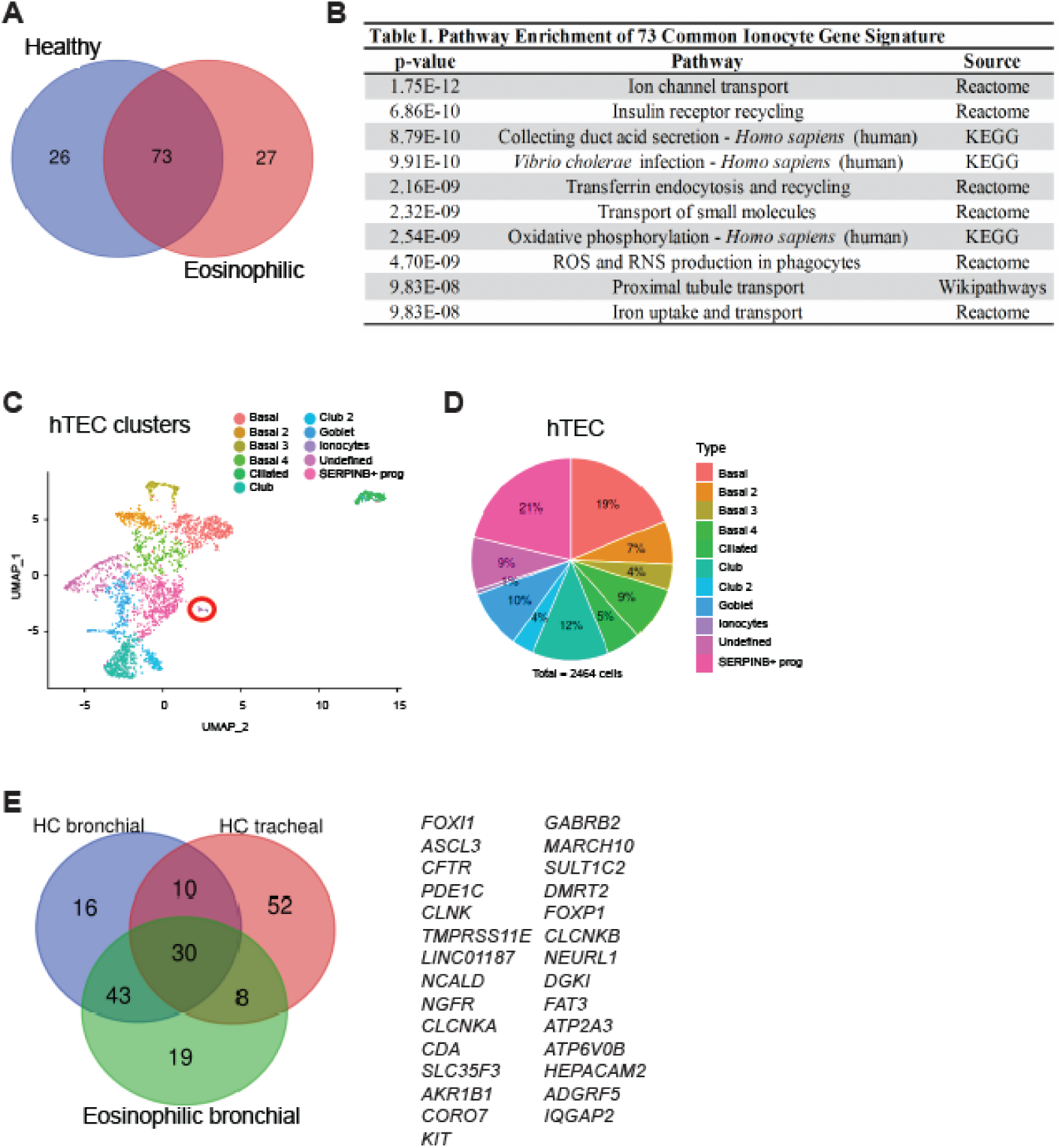
Pathway analysis of ionocyte gene signature and conservation throughout conducting airways. (A) Venn diagram of the 73 common ionocyte genes between hBECs from healthy subjects and eosinophilic asthma patients. (B) Pathway over-representation analysis using ConsensusPathDB of the 73 common ionocyte gene signature. (C) Human tracheal epithelial cells (hTECs) cultured at ALI from healthy donors (n= 4), were clustered by using a graph-based shared nearest neighbour method and plotted by UMAP. 11 clusters of cells were identified and then known cell types were classified using classical gene markers enriched in the clusters through leadingEdge. Transitory progenitors or non-classified cell types were identified by predominant transcriptional signatures SERPINB+ prog, or as Undefined. (D) The proportion of each cell type among hTECs (2464 cells). (E) Venn diagram of the conserved 30 common ionocyte gene signature among healthy hBECs, healthy hTECs, and hBECs from eosinophilic asthma. HC; healthy control.

### Quantification of ionocytes by protein markers confirms the loss of ionocytes in severe non-eosinophilic asthma hBECs

Using another independent cohort of patient hBECs differentiated at ALI we sort to validate the loss of ionocyte in non-eosinophilic asthma that we observed through scRNAseq. We quantified the ionocyte markers FOXI1, ASCL3, and CFTR by immunofluorescent protein quantification, which confirmed a significant almost complete loss of ionocytes (measured by dual staining ASCL3+FOXI1+ and CFTR+FOXI1+) in hBECs of non-eosinophilic asthma (Fig 5A-C). Although *CFTR* is found expressed to a lower level in other airway epithelial cell subtypes, in particular secretory cells, the highest expression of *CFTR* was in ionocytes from both healthy and asthma donors (Figure 3A). This is confirmed at the protein level where the overall number of CFTR-expressing cells was reduced in severe non-eosinophilic asthma (Fig 1E), which also validated the reduced CFTR function in hBECs from these patients (Fig 1A). We examined the impact on total CFTR protein-expressing cells by increasing secretory cell differentiation in the hBECs via recombinant IL-13 treatment. As expected this cytokine treatment increased secretory cells in the cultures, but it did not significantly increase CFTR expression (Supp Fig 6A-C). Similar to the scRNAseq results the immunofluorescent protein quantification showed there was a significant reduction of ionocytes in paucigranulocytic asthma hBECs and a complete loss in neutrophilic asthma (Supp Fig 6D-E). Established protein markers of other airway epithelial cell subsets were also quantified and showed no significant changes between healthy and non-eosinophilic asthma hBECs including acetylated tubulin+ ciliated cells, p63+ basal stem cells, or CCSP+ club cells, nor was there any functional difference in ciliated cell beat frequency between the patient hBECs (Fig 5D-E and Supp Fig 6F). Quantification of MUC5AC+ cells showed a significant increase in mature goblet cells in non-eosinophilic asthma (Fig 5D and Fig 2). Mucus metaplasia and hypersecretion are also features of other chronic airways disease with a dysfunctional CFTR pathway including CF and COPD. These results validated the findings from the scRNAseq discovery set analysis. Overall, these results clearly demonstrate a loss of ionocytes and increase in mature goblet cells in hBECs from non-eosinophilic asthma.

**Figure 5.**
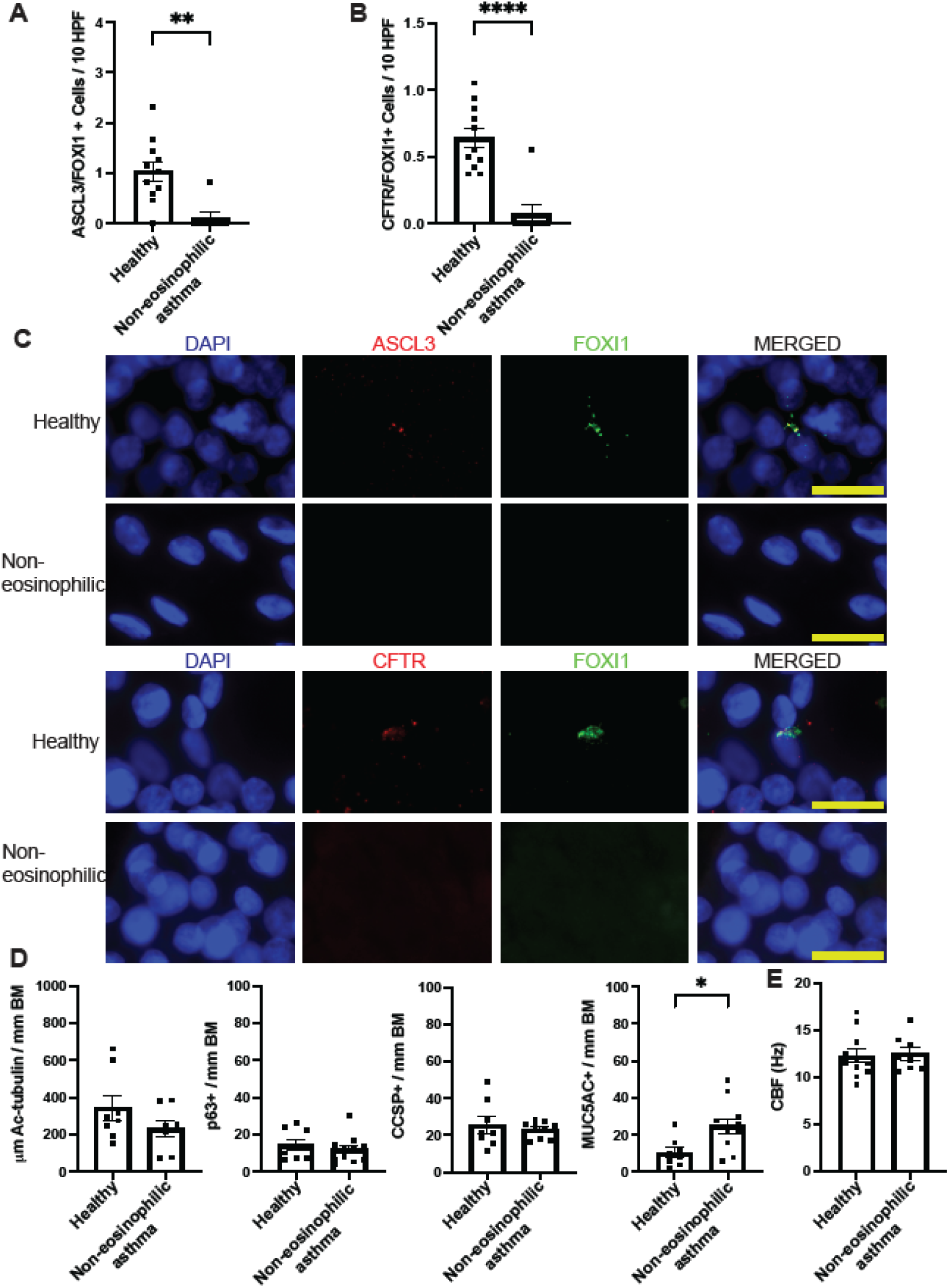
Quantification of ionocytes in hBECs by dual protein marker expression. Transwell membranes of ALI hBEC cultures from healthy or non-eosinophilic asthma were used for dual marker immunofluorescent quantification of ionocytes and other airway epithelial subtypes. (A) Dual ASCL3+FOXI1+ cells or (B) CFTR+FOXI1+ cells were quantified by immunofluorescence in hBECs from healthy donors or non-eosinophilic asthma, n=11. Values expressed per 10 high powered fields (HPF), mean ± SEM. (C) Representative immunofluorescent images for (A) and (B), scale bars represent 20 μm. (D) Quantification of common airway epithelial cell subtypes; ciliated (Ac-tubulin+), basal stem cells (p63+), club cells (CCSP+), and goblet (MUC5AC+) by immunofluorescence in hBECs from healthy donors (n=8) or non-eosinophilic asthma (n=8-11). Values expressed per mm of basement membrane (BM), mean ± SEM. (E) Cilia beat frequency (CBF) measured in live images from hBECs of healthy (n=11) versus non-eosinophilic asthma (n=8). Values expressed in Hz as mean ± SEM. Unpaired *t* test was used to determine statistical significance, *p<0.05, ** p<0.01, ****P<0.0001.

### Inflammation-induced loss of ionocytes mediated via neutrophilic asthma-associated cytokines, in particular IFN-γ

We hypothesized that the loss of CFTR-expressing ionocytes in non-eosinophilic asthma is likely an acquired phenotype mediated by inflammation. The characteristic cytokines of eosinophilic/Type 2 asthma include IL-4, -5, and -13. In contrast, severe non-eosinophilic asthma, and in particular neutrophilic asthma, is marked by elevated expression of a mix of Type 1 and Type 17 cytokines including IFN-γ, IL-17A, TNF-α, and IL-22 in patient airway tissue biopsies, sputum, and BAL (2, 16, 35, 36). To assess whether these neutrophilic asthma cytokines might lead to inflammation-induced loss of ionocytes, hBECs from healthy donors at ALI (which contain ionocytes) were treated with individual cytokines or a representative mixture (Type1+17 cytokine mix: IFN-γ, IL-17A, TNF-α, and IL-22). Immunofluorescent quantification by both ASCL3+FOXI1+ and CFTR+FOXI1+ measures both showed a dramatic reduction in ionocytes due to this cytokine mixture (Fig 6A-B). Treatment of these hBECs with individual cytokines showed this effect was primarily due to IFN-γ with a smaller partial effect of IL-17A, TNF-α, and IL-22 (Fig 6C-D). In contrast, there was no significant impact of IL-13 on ionocytes. This data demonstrates that inflammatory cytokines characteristically elevated in non-eosinophilic asthma airways can reduce the presence of ionocytes.

**Figure 6.**
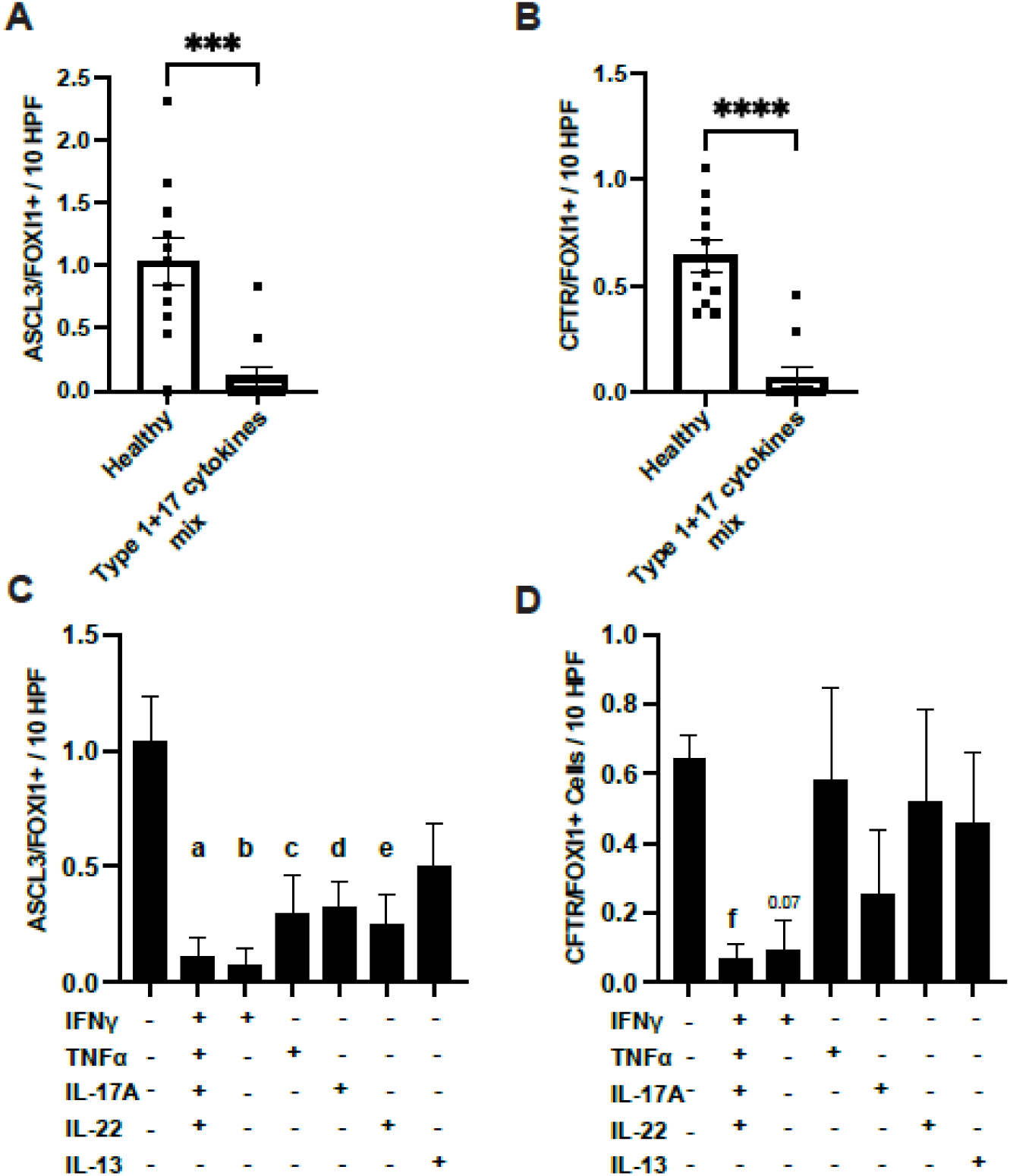
Inflammatory cytokine-induced loss of ionocytes. Transwell membranes of ALI hBEC cultures from healthy donors were used for dual marker immunofluorescent quantification of ionocytes. (A and C) Dual ASCL3+FOXI1+ cells or (B and D) CFTR+FOXI1+ cells were quantified by immunofluorescence in healthy hBECs treated with vehicle control, a cytokine mixture representative of non-eosinophilic asthma airway secretions including IFN-γ, IL-17A, IL-22, and TNF-α (n=11 each), or (C and D) with the individual cytokines indicated (n=7 each). Values expressed per 10 high powered fields (HPF), mean ± SEM. Paired *t* test (A, B) and One-way ANOVA with Dunnett’s multiple comparison test (C, D) were used to determine statistical significance. For (A and B) ***p<0.001, ****p<0.0001. For (C and D) a = p<0.0001 vs control, b = p<0.0001 vs control, c = p<0.01 vs control, d = p<0.01 vs control, e= p<0.01 vs control, f = p<0.05 vs control.

### Loss of ionocytes in a murine model of neutrophilic Type 2 low asthma but not eosinophilic Type 2 high asthma

Using LINCS L1000 Ligand Perturbations database and our ionocyte gene signature the *in silico* predictions suggested that IFN-γ was the most highly predicted molecule to regulate our ionocyte gene signature, which complemented the experimental findings from our *in vitro* results (Supp Fig 7A). To determine whether inflammation-induced loss of ionocytes also occurs *in vivo,* we quantified levels of ionocytes in established murine models of eosinophilic/Type 2 asthma versus neutrophilic asthma. We compared asthma models that are driven by the antigen ovalbumin (OVA) or the allergen fungus *Alternaria alternata* (AA), which are Type 2 high characterised by airway eosinophilia, versus a neutrophilic asthma model using the intracellular respiratory bacteria *Chlamydia muridarum* plus ovalbumin (Cmu/OVA) which produces a dominant airway neutrophilia (Fig 7A). These models are established to be driven by IL-4/-5/-13 or IFN-γ/IL-17A, respectively (37–40). We quantified ionocyte numbers in bronchial tissue measured by immunofluorescent quantification by both ASCL3+FOXI1+ and CFTR+FOXI1+. Ionocytes were significantly diminished in the Cmu/OVA neutrophilic asthma model, whereas these cells remained readily detectable in the eosinophilic models with either AA or OVA as well as PBS controls (Fig 7B-E). These *in vivo* data clearly support our *in vitro* studies with human airway epithelial cells demonstrating that neutrophilic asthma associated inflammation can induce loss of CFTR-expressing ionocytes.

**Figure 7.**
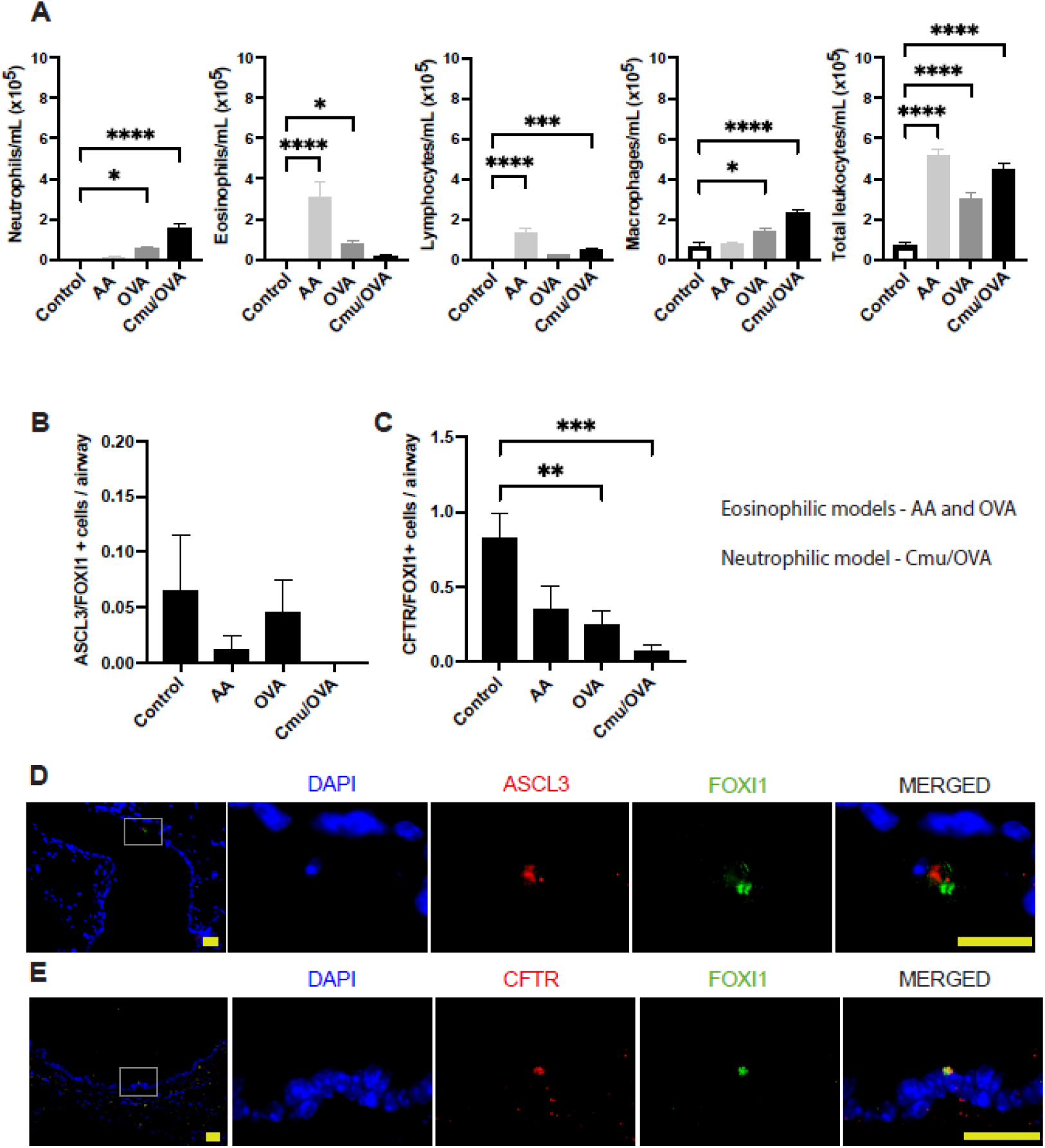
Loss of ionocytes *in vivo* in a murine model of neutrophilic asthma. (A) Number of neutrophils, eosinophils, lymphocytes, macrophages, and total number of cells/mL in bronchoalveolar lavage (BAL) fluid in control, fungal allergen *Alternaria alternata* (AA), and protein antigen ovalbumin (OVA) models of T2 asthma, as well as the *Chlamydia muridarum* (Cmu)/OVA-treated model of non-T2 asthma. (B) Dual ASCL3+FOXI1+ cells or (C) CFTR+FOXI1+ cells were quantified by immunofluorescence in the bronchi of mice from these asthma models (n=7-11 mice per group). Values expressed as per airway, mean ± SEM. (D and E) Representative immunofluorescent images of PBS control mouse airways stained with CFTR plus FOXI1 or ACSL3 plus FOXI1, scale bar is 20 µm. One-way ANOVA with Dunnett’s multiple comparison test compared to control mice, *p<0.05, **p < 0.01, ***p < 0.001, ****p < 0.0001.

## Discussion

Severe asthma (approximately 1 in 10 adult patients with asthma) is refractory to conventional therapies including inhaled corticosteroids and results in frequent hospitalizations as one of the largest disease burdens in the respiratory clinic (41, 42). Although the eosinophilic and Type 2 dominant subtype of severe asthma can now be treated successfully in many cases with monoclonal antibody therapies targeting IL-5/IL-5R, IgE, and IL-4Rα (IL-4/13) pathways, many severe asthma patients, in particular those with non-eosinophilic or dominant neutrophilic airway disease, remain poorly treated (1, 43). There is a clear need for new and effective therapies to target this patient subpopulation, which requires a deeper understanding of the cell and molecular mediators driving this subtype of asthma.

Insights into the pathogenesis of non-eosinophilic asthma may be gained by examining the crossover with cellular and pathological markers of other chronic airways diseases including COPD, CF, and subsets of non-CF bronchiectasis. These diseases also show strong neutrophilic infiltrates in the airways, a dominant mixture of Type 1 and Type 17 cytokine responses, mucous hypersecretion, and high airway bacterial colonization (4–9, 44). Indeed, the characteristic airway colonizers of CF, *Pseudomonas aeruginosa* and *Haemophilus influenzae,* are also found in high abundance in COPD and non-CF bronchiectasis and are microbiome markers of non-eosinophilic asthma (45–49). These similarities raise the possibility of common mechanistic drivers. A defective CFTR is sufficient to drive these disease features in CF, and is also linked to this phenotype in a proportion of non-CF bronchiectasis cases (50). Smoking is known to reduce the function of CFTR resulting in an acquired loss of CFTR activity in COPD (51, 52). A small case series report has also linked non-allergic asthma to abnormal CFTR function observed through nasal potential differences (12), providing a clue that defects in this pathway could be linked to the clinical phenotype of non-eosinophilic asthma.

Any defects in CFTR activity in non-eosinophilic asthma are likely acquired or induced rather than genetic. Although there is evidence, some of it conflicting, that the risk of asthma is higher in non-CF individuals with CFTR gene variants (i.e. CFTR carrier heterozygosity), the odds ratio suggests that genetic variants are unlikely to be a dominant factor in non-eosinophilic asthma (10, 11). Based on this rationale we investigated the function of the CFTR in airway epithelial cells from patients with non-eosinophilic asthma and observed a substantial reduction (∼50% loss) in CFTR function and protein expression. This confirmed our hypothesis that the CFTR pathway may be dysfunctional in airway epithelial cells from severe non-eosinophilic asthma. To provide greater mechanistic depth to this observation we used scRNAseq to analyse hBECs from severe asthma subtypes and healthy controls. scRNAseq is able to pick up transcriptional changes at the individual cell level and so for rarer cell types can be studied that would be missed with the more traditional transcriptional approaches of bulk RNAseq, microarrays, or targeted PCR. We discovered through scRNAseq that the CFTR-rich cell type, the airway ionocyte, was diminished in hBECs from non-eosinophilic asthma. More specifically this cell types was almost completely lost in neutrophilic asthma and significantly diminished in paucigranulocytic asthma, which we were able to confirm using immunofluorescent techniques for dual staining and quantification of known ionocyte markers. This was a unique observation as both scRNAseq and confirmatory protein cell marker validation showed that other epithelial subtypes remain largely unchanged with the exception of a greater number of mature goblet cells in neutrophilic asthma observed both in the scRNAseq data and immunofluorescent quantification of MUC5AC+ cells. As goblet cell metaplasia and mucus hypersecretion are observed in other chronic airways diseases linked to CFTR dysfunction (CF and COPD) this provided further evidence of a role in non-eosinophilic asthma.

Using the cell cluster marker genes we identified in ionocytes in hBECs from healthy controls and eosinophilic asthma as well as those identified in healthy hTECs we discovered a conserved ionocyte gene signature present in the trachea to the 3^rd^-4^th^ branch of the bronchi. This was a markedly conserved gene signature (∼30%) from these 3 different sets of donors. This signature confirmed previously known ionocyte markers (53, 54) like the transcription factor *FOXI1*, *ACSL3*, *CFTR*, *PDE1C*, and *ATP6V0B* but also identified new markers with potential interesting cellular biology relevant to ionocyte function including *NEURL1*, a Notch pathway regulator, and *NCALD*, *GABRB2*, and *NGFR* all involved in neural signalling and neurotransmission. Future studies will be required to investigate the interaction of ionocytes and neural signalling processes in the airways. Overall pathway analysis clearly showed that the ionocyte gene signature was linked to ion channel transport for chloride, iron, and potassium and pH regulation. It can be speculated from these results that ionocytes may regulate ion homeostasis and pH at the airway surface at least in part through neurotransmission pathways. This would suggest these cells could be important in a range of chronic airways diseases where ion homeostasis and pH are dysregulated. Indeed, asthma patients with lower lung function and worse symptoms have lower airway pH detected in exhaled breath condensate and dysregulated iron homeostasis (55, 56).

Our scRNAseq analysis of hBECs also identified two subpopulations of clusters of epithelial cells (we referred to as *LAMB3+* or *SERPINB+* progenitors) without a clear delineated differentiated cell identity and are likely transitioning between subtypes. These clusters expressed a limited number of markers in common with basal cells, such as *SOX* gene family members, cyclin cell cycle genes, and in the case of one of the populations the basal cell gene *LAMB3*, suggesting they may have progenitor potential. These subpopulations also had consistent expression across the patient subgroups of either *LAMB3* or *SERPINB* family genes, hence the nomenclature we used, and their percentages was very similar across all patient groups. It is likely these clusters represent a transitory or plastic stage of differentiation of the airway epithelial cells. The *LAMB3+* progenitor cluster had a signature enriched for some epithelial-mesenchymal genes and the *SERPINB+* progenitor cluster had genes involved in protease-anti-protease pathways. Future studies will need to confirm these subpopulations *in vitro* and *in vivo* and investigate their cellular function.

Our data demonstrated that ionocytes are lost in hBEC ALI cultures from non-eosinophilic asthma patients and we hypothesized that this is likely an acquired or induced defect caused by inflammatory pathways. Differentiated ALI cultures of airway epithelial cells are well established to retain memory of many of the features of human patients *in vivo* retaining the properties of p63+ basal stem cell hyper-proliferation, diminished interferon and anti-viral responses and increased mucus cell metaplasia observed in asthma and COPD (57–61). ALI airway epithelial cultures also reflect global gene expression changes of airway epithelial brushings taken from patients (62, 63). Thus, we postulated an inflammation-induced loss of ionocytes and CFTR function was responsible for the reduction observed in hBECs from non-eosinophilic asthma patients. This would constitute an ‘inflammatory memory’ as proposed by *Ordovas-Montanes e al.,* being programmed into the epithelial progenitor cells (64). The airways of non-eosinophilic asthma patients, in particular the neutrophilic subset, are consistently found to be dominated by Th1 and Th17 cytokines in BAL, endobronchial biopsies, and sputum with elevated levels of IL-17A, IL-22, IFN-γ, and TNF-α (2, 16, 35). Based on this knowledge we treated hBECs from healthy non-asthmatic donors with these cytokines in a cocktail, or individually, and demonstrated that these mediators of neutrophilic airway inflammation are sufficient to dramatically reduce the numbers of CFTR-expressing ionocytes. In particular, the Type 1 cytokine IFN-γ, whose protein levels and downstream gene signatures are enriched in neutrophilic asthma (2, 16, 35) almost completely inhibited ionocyte differentiation.

To confirm these *in vitro* results we used multiple mouse models of asthma-like airway pathology that recapitulate either a classical Type 2 cytokine/eosinophilic-dominant airways disease (through sensitisation and challenge with ovalbumin (OVA) or *Alternaria alternata* (AA)), or a more neutrophilic-dominant airways disease characterised by elevated Th1 and Th17 cytokines induced by *Chlamydia muridarum* infection during OVA challenge (37–40). Our *in vivo* findings from these models also showed an almost complete loss of airway ionocytes in the presence of the neutrophilic airway disease. These studies add further weight to our observations that inflammation-induced loss of ionocytes and CFTR is a feature of non-eosinophilic asthma. Future studies will be necessary to determine whether ionocytes have a role in mediating airway ion transport homeostasis and regulating airway pH, other than through the CFTR, and how this impacts asthma pathogenesis in these models.

This work highlights that an important pathogenic ion transport pathway through the CFTR and rarer cell type, the ionocyte, are both dysregulated in severe non-eosinophilic asthma. It also demonstrates the tremendous value of scRNAseq in discovering changes in rarer cell types that are otherwise missed by the averaging effect of bulk sample analysis. Given the need for more effective treatment options for this patient population, and the role of this pathway in other chronic airway diseases, this may represent a novel therapeutic target for precision medicine guided treatment of severe asthma.

## Acknowledgements

LC and GAH conducted the experimental studies and analysed data, PP conducted experimental work, AR, CK, and THY contributed to data analysis, SB, RK, PH provided critical reagents/samples, PSF helped draft manuscript, JH provided critical samples and analysis, CR conducted data analysis, PW and GK designed studies, analysed data, and wrote the manuscript.

---

**Supp figure 1.** Characterisation of patients with severe asthma subtypes and single-cell RNA-seq analysis. (A) Percentage from differential cell counts in BAL fluid from healthy subjects and asthma patients subtyped. Single-cell RNA-seq was performed on single-cell suspensions generated from hBECs of eosinophilic asthma patients and paucigranulocytic asthma patients (n=4 each). (B) Cells were clustered by using a graph-based shared nearest neighbour method and plotted by UMAP together with heatmaps of gene UMI counts. 9 clusters of cells were identified in all samples and then known cell types were classified using classical gene markers enriched in the clusters through leadingEdge. Transitory progenitors or non-classified cell types were labelled by predominant transcriptional signatures SERPINB+ prog, LAMB3+ prog, or Undefined. (C) The proportion of each cell type in eosinophilic (9624 cells) and paucigranulocytic asthma (6914 cells) patients were calculated using pie charts.

**Supp figure 2.** Unsupervised hierarchical clustering of all patient samples and cell subtypes (columns) by known ionocyte and basal cell marker genes (rows), visualised using a heatmap plot.

**Supp figure 3.** (A) Violin plots of expression of ionocyte gene markers (*PDE1C, FOXI1, ASLC3* and *CFTR*) in each of the 9 cell subtype clusters from eosinophilic versus paucigranulocytic asthma patients.

**Supp figure 4.** Weighted dot plot showing top 10 features of each cluster from the scRNAseq data derived from cells of (A) neutrophilic, (B) eosinophilic, and (C) paucigranulocytic asthma patients. Each dot is sized to represent the percent of cells in each cluster expressing the corresponding top 10 gene, and colors represent the average expression of each maker gene across within that cluster.

**Supp figure 5.** Pathway over-representation analysis. (A) Gene Ontology (GO) analysis for Biological Process and (B) Molecular Process of the 73 common ionocyte genes. (C) Human Phenotype Ontology analysis of the 73 common ionocyte genes. (D) Weighted dot plot showing top 10 features of each cluster from the scRNAseq data derived from human tracheal epithelial cells (hTECs) from healthy donors. Each dot is sized to represent the percent of cells in each cluster expressing the corresponding top 10 gene, and colors represent the average expression of each maker gene across within that cluster. (E) Pathway over-representation analysis using ConsensusPathDB of the 30 common genes in the ionocyte gene signature from Fig 4E conserved between hBECs of healthy and eosinophilic asthma donors as well as hTECs.

**Supp figure 6.** Loss of ionocytes in non-eosinophilic (neutrophilic and paucigranulocytic) asthma hBECs. (A) *CFTR* gene expression, and (B) quantification of the number of CFTR protein expressing (CFTR+) cells in healthy hBECs treated with or without IL-13. (C) Representative immunofluorescent images of healthy hBECs treated with or without IL-13 showing increased numbers of CCSP+ and MUC5AC+ goblet cells. (D) ASCL3+FOXI1+ and (E) CFTR+FOXI1+ in hBECs from neutrophilic and paucigranulocytic asthma or healthy donors (n=4-11), values expressed as mean ± SEM. (F) Representative immunofluorescent images of healthy hBECs indicating cells positive for MUC5AC, Ac-tubulin, p63 (green), or CCSP (red), scale bar is 20μm. *p ≤ 0.05, **p < 0.01, ***p < 0.001.

**Supp figure 7.** (A) Library of Integrated Network-Based Cellular Signatures (LINCS) L1000 ligand perturbations analysis showing cytokines and ligands most highly predicted to regulate the ionocyte gene signature.

