## Supplemental figures for "Single cell RNA-seq identifies inflammation-induced loss of CFTR-expressing airway ionocytes in non-eosinophilic asthma"

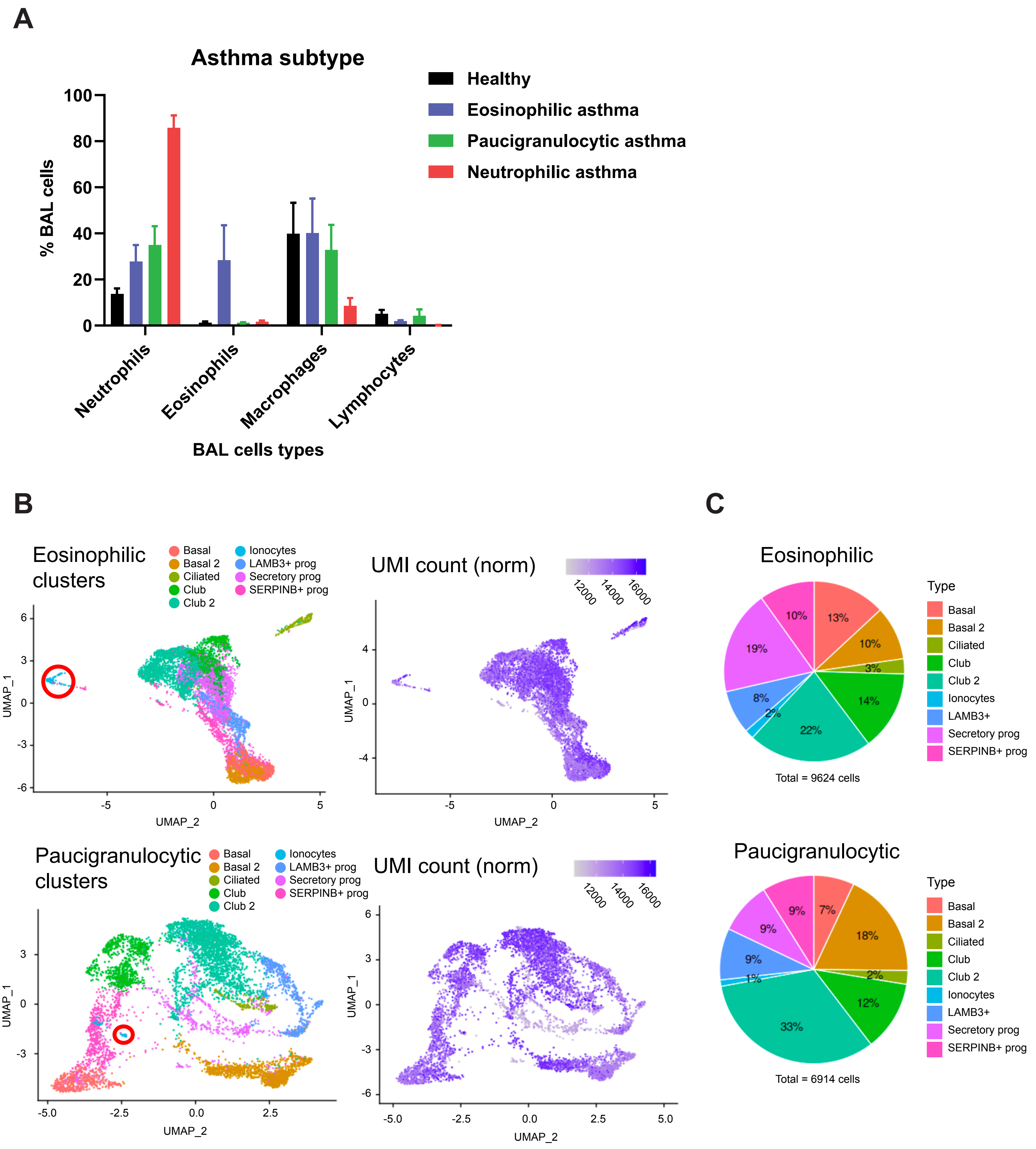

A
Asthma subtype
Healthy
Eosinophilic asthma
Paucigranulocytic asthma
Neutrophilic asthma
% BAL cells
100
80
60
40
20
0
Neutrophils
Eosinophils
Macrophages
Lymphocytes
BAL cells types
B
Eosinophilic clusters
Basal
Basal 2
Ciliated
Club
Club 2
Ionocytes
LAMB3+ prog
Secretory prog
SERPINB+ prog
UMAP_1
UMAP_2
UMI count (norm)
12000
14000
16000
Paucigranulocytic clusters
C
Eosinophilic
Type
Basal
Basal 2
Ciliated
Club
Club 2
Ionocytes
LAMB3+
Secretory prog
SERPINB+ prog
13%
10%
3%
14%
22%
2%
8%
19%
10%
Total = 9624 cells
Paucigranulocytic
7%
18%
2%
12%
33%
1%
9%
9%
9%
Total = 6914 cells

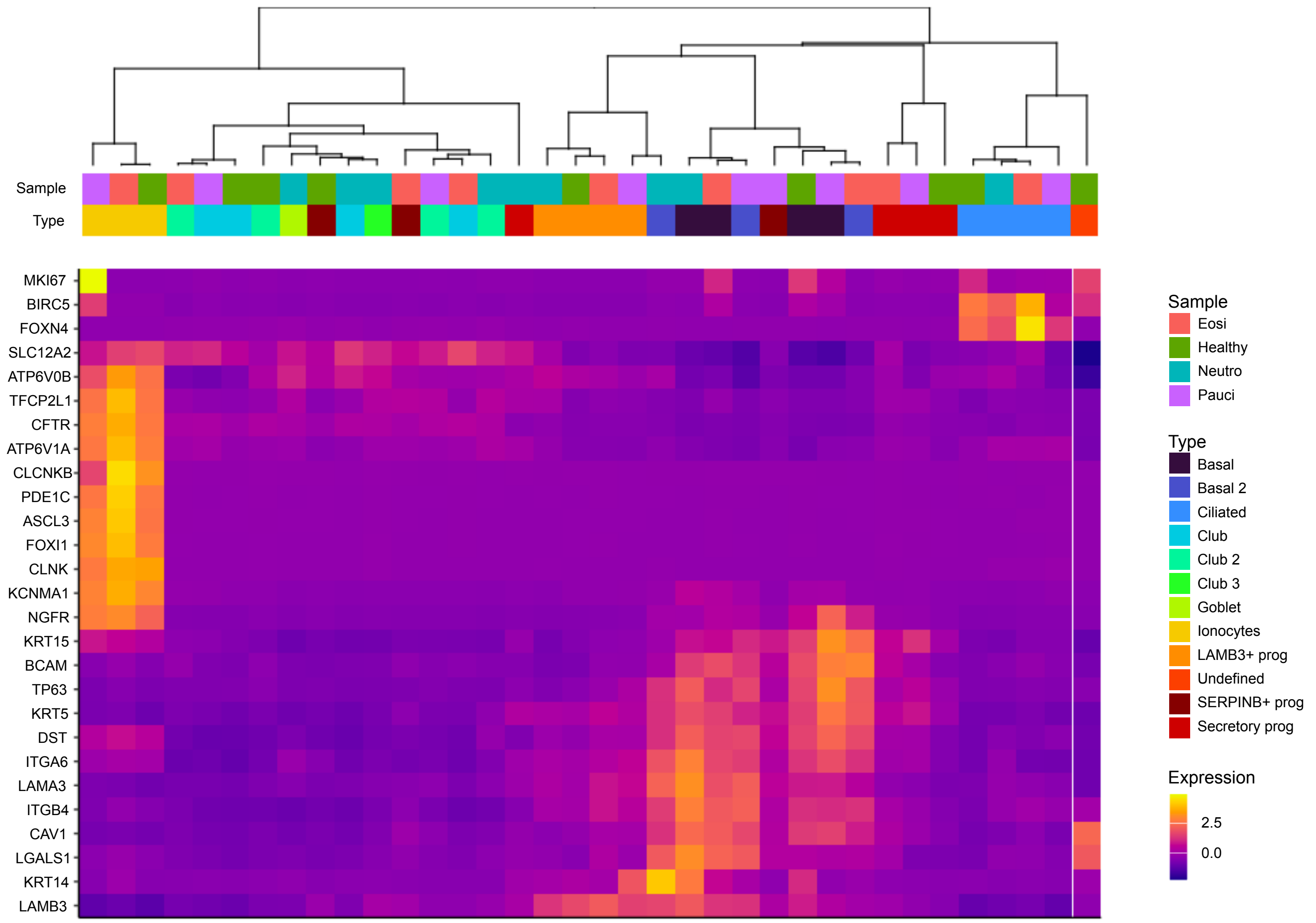

Sample
Type
MKI67
BIRC5
FOXN4
SLC12A2
ATP6V0B
TFCP2L1
CFTR
ATP6V1A
CLCNKB
PDE1C
ASCL3
FOXI1
CLNK
KCNMA1
NGFR
KRT15
BCAM
TP63
KRT5
DST
ITGA6
LAMA3
ITGB4
CAV1
LGALS1
KRT14
LAMB3
Sample
Eosi
Healthy
Neutro
Pauci
Type
Basal
Basal 2
Ciliated
Club
Club 2
Club 3
Goblet
Ionocytes
LAMB3+ prog
Undefined
SERPINB+ prog
Secretory prog
Expression
2.5
0.0

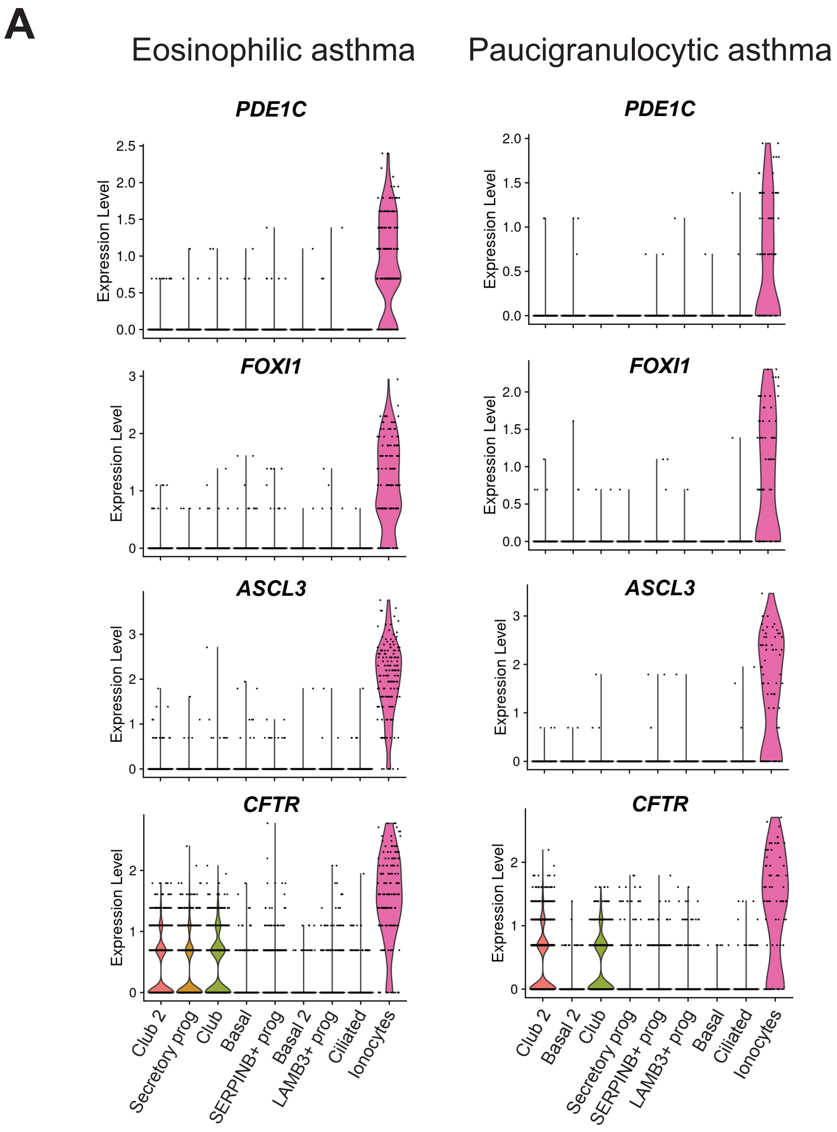

A
Eosinophilic asthma
Paucigranulocytic asthma
PDE1C
FOXI1
ASCL3
CFTR
Expression Level
Club 2
Secretory prog
Club
Basal
SERPINB+ prog
Basal 2
LAMB3+ prog
Ciliated
Ionocytes

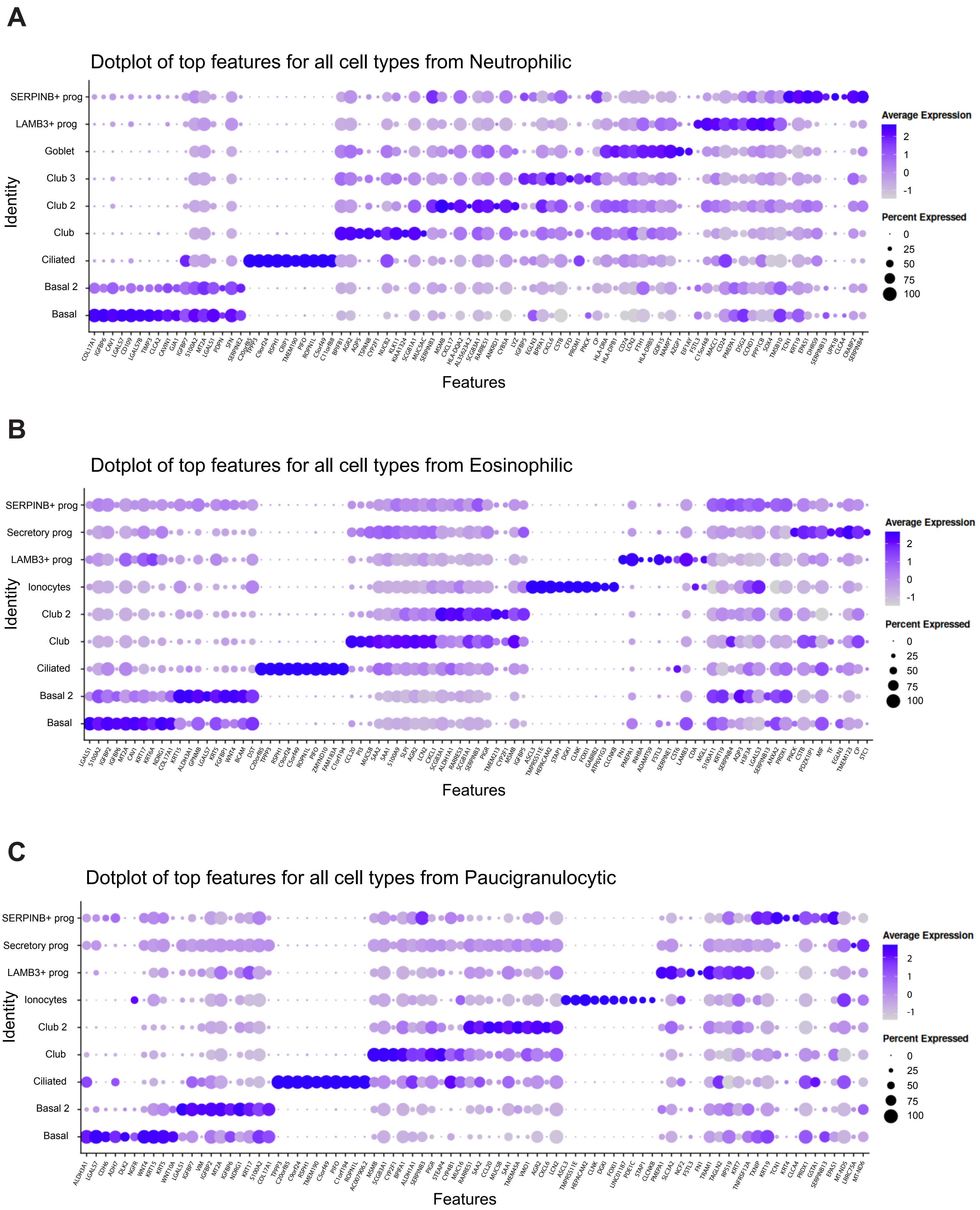
A
Dotplot of top features for all cell types from Neutrophilic
SERPINB+ prog
LAMB3+ prog
Goblet
Club 3
Club 2
Club
Ciliated
Basal 2
Basal
Identity
Features
Average Expression
2
1
0
-1
Percent Expressed
0
25
50
75
100
B
Dotplot of top features for all cell types from Eosinophilic
SERPINB+ prog
Secretory prog
LAMB3+ prog
Ionocytes
Club 2
Club
Ciliated
Basal 2
Basal
Identity
Features
Average Expression
2
1
0
-1
Percent Expressed
0
25
50
75
100
C
Dotplot of top features for all cell types from Paucigranulocytic
SERPINB+ prog
Secretory prog
LAMB3+ prog
Ionocytes
Club 2
Club
Ciliated
Basal 2
Basal
Identity
Features
Average Expression
2
1
0
-1
Percent Expressed
0
25
50
75
100

**A**

| Table II. GO Biological Process | | |
|---|---|---|
| **Index** | **Name** | **Adjusted p-value** |
| 1 | Phagosome acidification (GO:0090383) | 4.66E-07 |
| 2 | Intracellular pH reduction (GO:0051452) | 7.01E-07 |
| 3 | Transferrin transport (GO:0033572) | 7.01E-07 |
| 4 | Phagosome maturation (GO:0090382) | 7.01E-07 |
| 5 | Positive regulation of B cell receptor signaling pathway (GO:0050861) | 0.01291 |
| 6 | Iron ion transport (GO:0006826) | 3.69E-06 |
| 7 | Bone cell development (GO:0098751) | 0.01996 |
| 8 | Regulation of receptor binding (GO:1900120) | 0.01996 |
| 9 | Negative regulation of receptor binding (GO:1900121) | 0.01996 |
| 10 | Proton transmembrane transport (GO:1902600) | 0.000521 |

**B**

| Table III. GO Molecular Process | | |
|---|---|---|
| **Index** | **Name** | **Adjusted p-value** |
| 1 | ATPase activity, coupled to transmembrane movement of irons, rotational mechanism (GO:0044769) | 1.12E-07 |
| 2 | Proton-transporting ATPase activity, rotational mechanism (GO:0046961) | 1.12E-07 |
| 3 | Pyrophosphate hydrolysis-driven proton transmembrane transporter activity (GO:0009678) | 2.60E-08 |
| 4 | Voltage-gated chloride channel activity (GO:0005247) | 1.36E-02 |
| 5 | Chloride channel activity (GO:0005254) | 0.000128 |
| 6 | Chloride channel regulator activity (GO:0017081) | 2.69E-02 |
| 7 | Calcitonin family receptor activity (GO:0097642) | 0.1393 |
| 8 | Uridine transmembrane transporter activity (GO:0015213) | 0.1393 |
| 9 | Purine nucleotide transmembrane transporter activity (GO:0015216) | 0.1393 |
| 10 | Nerve growth factor binding (GO:0048046) | 0.1393 |

**C**

| Table IV. Human Phenotype Ontology | | |
|---|---|---|
| **Index** | **Name** | **Adjusted p-value** |
| 1 | Abnormality of chloride homeostasis (HP:0011422) | 1.97E-07 |
| 2 | Metabolic alkalosis (HP:0200114) | 1.51E-06 |
| 3 | Hypokalemic alkalosis (HP:0001949) | 1.89E-06 |
| 4 | Alkalosis (HP:0001948) | 9.81E-07 |
| 5 | Hyperaldosteronism (HP:0000859) | 0.000005529 |
| 6 | Renal salt wasting (HP:0000127) | 6.43E-06 |
| 7 | Hypokalemia (HP:0002900) | 0.000001354 |
| 8 | Hyponatremia (HP:0002902) | 0.00001506 |
| 9 | Abnormality of potassium homeostasis (HP:0011042) | 0.000005012 |
| 10 | Hyperactive renin-angiotensin system (HP:0000841) | 0.003607 |

**D**

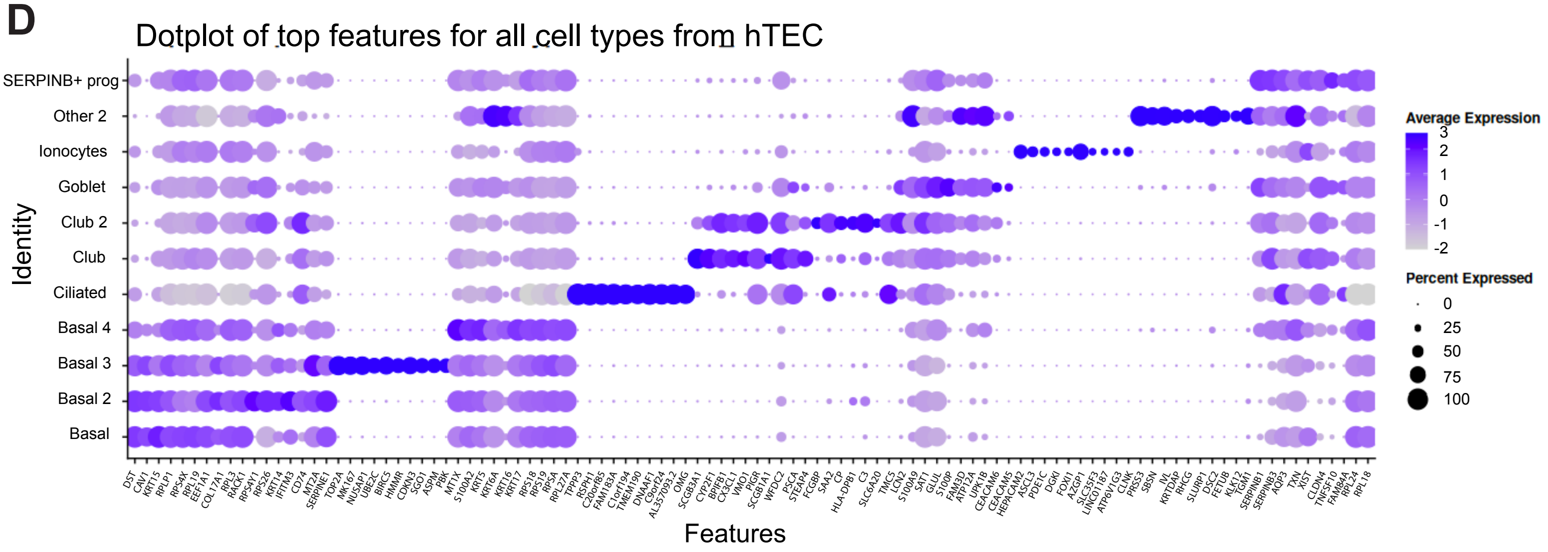


**E**

| Table V. Pathway Enrichment of 30 Common Ionocyte Gene Signature | | |
|---|---|---|
| **p-value** | **Pathway** | **Source** |
| 5.87E-05 | Ion channel transport | Reactome |
| 1.16E-03 | mBDNF and proBDNF regulation of GABA neurotransmission | Wikipathways |
| 1.24E-03 | Transport of small molecules | Reactome |
| 2.00E-03 | *Vibrio cholerae* infection - *Homo sapiens* (human) | KEGG |
| 2.06E-03 | Signal transduction | Reactome |
| 2.96E-03 | Glycerolipid metabolism - *Homo sapiens* (human) | KEGG |
| 6.47E-03 | Morphine addiction - *Homo sapiens* (human) | KEGG |
| 6.89E-03 | Stimuli-sensing channels | Reactome |
| 7.47E-03 | Pathways regulating hippo signaling | Wikipathways |
| 8.07E-03 | Pancreatic secretion - *Homo sapiens* (human) | KEGG |

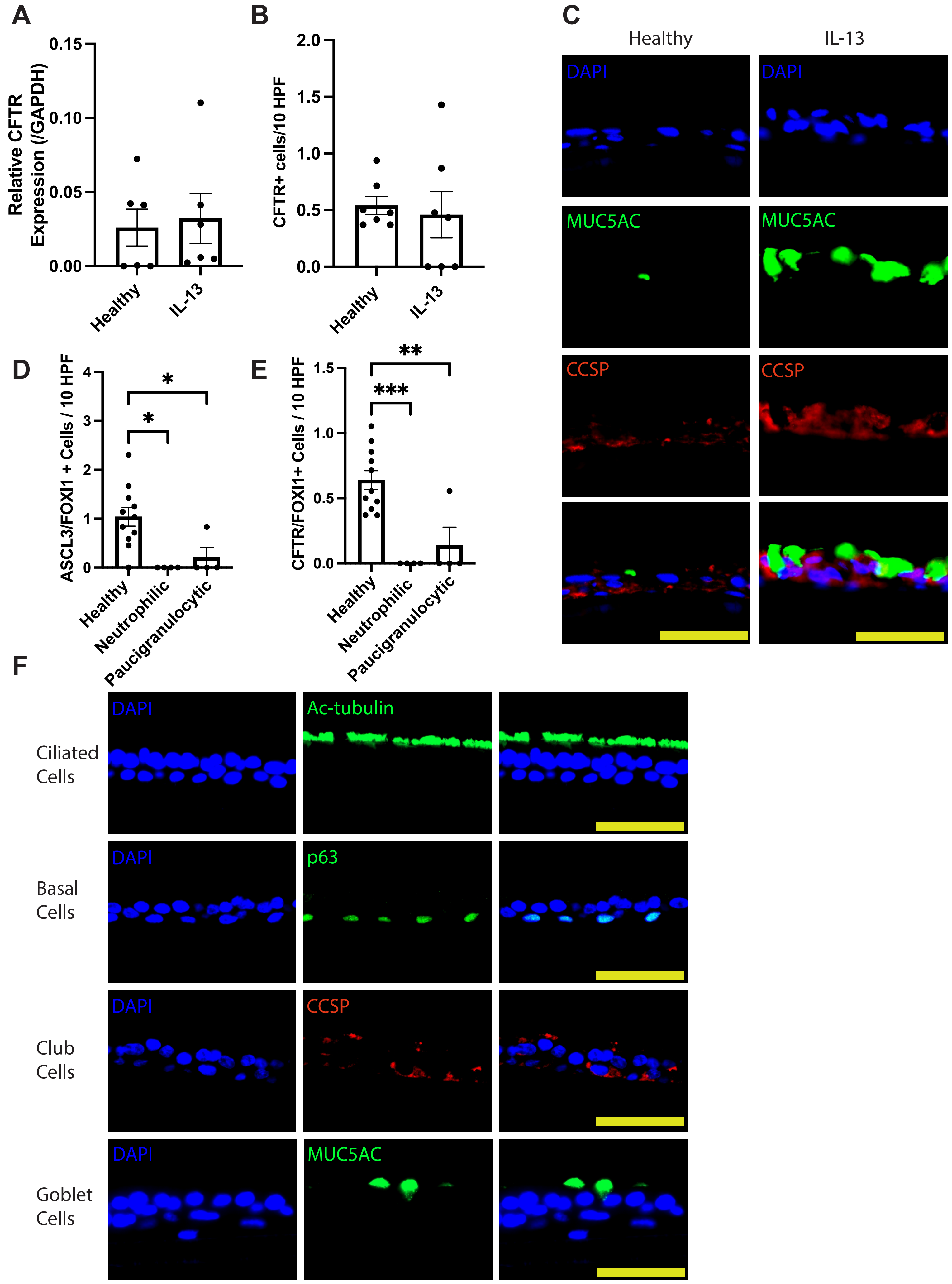
A
Relative CFTR Expression (/GAPDH)
0.15
0.10
0.05
0.00
Healthy
IL-13
B
CFTR+ cells/10 HPF
2.0
1.5
1.0
0.5
0.0
Healthy
IL-13
C
Healthy
IL-13
DAPI
MUC5AC
CCSP
D
ASCL3/FOXI1 + Cells / 10 HPF
4
3
2
1
0
*
*
Healthy
Neutrophilic
Paucigranulocytic
E
CFTR/FOXI1+ Cells / 10 HPF
1.5
1.0
0.5
0.0
**
***
Healthy
Neutrophilic
Paucigranulocytic
F
Ciliated Cells
DAPI
Ac-tubulin
Basal Cells
DAPI
p63
Club Cells
DAPI
CCSP
Goblet Cells
DAPI
MUC5AC

**A**

| Table VI. LINCS L1000 Ligand Perturbations | | |
|---|---|---|
| **Index** | **Name** | **Adjusted p-value** |
| 1 | IFNG-MDAMB231 | 3.02E-04 |
| 2 | MCSF-5KBR3 | 1.08E-03 |
| 3 | IFNG-MCF7 | 9.18E-04 |
| 4 | IL6-MCF7 | 1.30E-03 |
| 5 | IFNG-MCF10A | 0.001488 |
| 6 | BNGF-H5578T | 4.28E-03 |
| 7 | IL1-MDAMB231 | 0.004282 |
| 8 | IFNA-MCF7 | 0.004282 |
| 9 | BTC-BT20 | 0.004282 |
| 10 | SCF-SKBR3 | 0.004282 |
